# Anticipatory and evoked visual cortical dynamics of voluntary temporal attention

**DOI:** 10.1101/2022.11.18.517084

**Authors:** Rachel N. Denison, Karen J. Tian, David J. Heeger, Marisa Carrasco

## Abstract

We can often anticipate the precise moment when a stimulus will be relevant for our behavioral goals. Voluntary temporal attention, the prioritization of sensory information at task-relevant time points, enhances visual perception. However, the neural mechanisms of voluntary temporal attention have not been isolated from those of temporal expectation, which reflects timing predictability rather than relevance. Here we used time-resolved steady-state visual evoked responses (SSVER) to investigate how temporal attention dynamically modulates visual activity when temporal expectation is controlled. We recorded MEG while observers directed temporal attention to one of two sequential grating targets with predictable timing. Meanwhile, we used a co-localized SSVER probe to continuously track visual cortical modulations both before and after the target stimuli. In the pre-target period, the SSVER gradually ramped up as the targets approached, reflecting temporal expectation. Furthermore, we found a low-frequency modulation of the SSVER, which shifted approximately half a cycle in phase according to which target was attended. In the post-target period, temporal attention to the first target transiently modulated the SSVER shortly after target onset. Thus, temporal attention dynamically modulates visual cortical responses via both periodic pre-target and transient post-target mechanisms to prioritize sensory information at precise moments.

## Introduction

We can often anticipate the points in time at which sensory events will be relevant for our behavior. For example, when playing table tennis, it is critical to see the ball at the moment it bounces on the table; only seeing the ball half a second earlier or later would be unhelpful (1). Voluntary temporal attention is the deliberate prioritization of a point in time that we know in advance will be relevant for our behavioral goals (2).

Voluntary temporal attention involves prioritizing a stimulus on the basis of both its timing and its relevance to the observer’s task (3, 4). These two features distinguish voluntary temporal attention from other dynamic phenomena (5). For example, a task-relevant stimulus may appear at an unpredictable time, eliciting reactive attentional dynamics—as in the attentional blink (6–8)— rather than the proactive dynamics of voluntary temporal attention. Alternatively, an irrelevant stimulus may appear at a predictable time, eliciting temporal expectation but not attentional prioritization (5, 9–11).

Voluntary temporal attention has been shown to improve behavioral performance (3, 4, 12, 13) and affect microsaccades (10, 14) over and above the effects of temporal expectation, suggesting dissociable neural mechanisms. Likewise, in the domains of visual spatial and feature-based attention, as well as auditory temporal attention, relevance and predictability can have distinct behavioral and neural effects (9, 15–24). Finally, spatial (25, 26) and feature-based (27) attention have larger effects on visual responses to targets that appear at expected vs. unexpected times, further suggesting that temporal attention and expectation may have separable effects on visual cortex.

The neural mechanisms specific to voluntary temporal attention are unclear, however, because previous neural studies have typically manipulated visual temporal attention and temporal expectation together, by presenting a single task-relevant target at a more or less predictable time (28–33). As a result, the neural modulations observed could be related to timing predictability, time point relevance, or a combination of the two. Such studies have found increased visual responses to temporally anticipated targets (31, 33) as well as slow ramping activity in dorsal stream cortical and subcortical areas (34–37) that peaked around the time of an anticipated event. These ramping signals could be related to time-keeping and top-down control of anticipatory processes, including motor preparation. Temporal expectation has also been associated with neural oscillations in sensory and fronto-parietal cortices, particularly in the delta (0.5-4 Hz) range, both during rhythmic stimulation at the delta frequency (38–40) and in predictable interval timing tasks (41).

Our normalization model of dynamic attention, which was fit to psychophysical data (3, 4, 13), predicts that voluntary temporal attention will change the gain of visual responses rapidly and transiently, roughly coincident with the evoked sensory response to the visual stimulus (3), rather than slowly and gradually, before the stimulus appears. Importantly, sensory gain modulations that multiplicatively scale neural response amplitudes can only be measured in the presence of a probe stimulus that elicits non-zero visual activity. A few studies of temporal expectation presented probe stimuli during a pre-target period and found anticipatory modulation of sensory activity, including ramping up of gamma band signals in V1 (42), ramping of responses to auditory probe stimuli in A1 (43), and spatial attentional modulation that depended on task timing in V4 (26), consistent with anticipatory gain modulation (3, 44). However, it remains unclear whether voluntary temporal attention still modulates visual gain when expectation is controlled and what the dynamics of any such modulation may be.

Here we investigated how temporal attention dynamically modulates visual cortical responses using MEG. Human observers performed a challenging orientation discrimination task in which voluntary temporal attention was manipulated independent of temporal expectation, which was constant. A precue directed temporal attention to one of two brief, sequential grating targets that were presented at fixed, predictable times. To measure attentional modulation continuously across time, we used time-resolved steady-state visual evoked responses (SSVER) (45–50). The grating targets were superimposed on a 20 Hz flickering noise probe, which generated a continuous SSVER signal (51, 52). We tracked this signal throughout the trial—both leading up to and following the target stimuli—by calculating the inter-trial phase coherence (ITPC) at the stimulus frequency in a time-resolved fashion. ITPC is the primary driver of SSVER power (51–54) and indexes the reliability with which the visual stimulus drives cortical responses: a more reliable neural response to the stimulus will yield a higher phase coherence across trials (51, 55).

We found dynamic task-related modulations of visual cortical responses leading up to and following target stimuli. Visual cortical ITPC slowly ramped up in anticipation of the predictable target stimuli regardless of the precue, an overall effect of temporal expectation. Furthermore, we found a low-frequency (2 Hz) modulation of the ITPC in the pre-target period, which shifted approximately 180 degrees in phase according to which target was attended. In the post-target period, temporal attention to the first target transiently modulated ITPC shortly after target onset, increasing the peak response to the target when it was attended. The findings show how both pre- and post-target modulatory mechanisms contribute to voluntary temporal attention. The presence and timecourse of these dynamic modulations in visually-driven cortical activity suggest mechanisms for how temporal attention can selectively enhance visual perceptual sensitivity at specific moments in time.

## Results

### Temporal precueing improved perceptual sensitivity

To investigate the effects of voluntary temporal attention on visual cortical dynamics, we recorded MEG while observers performed a temporal attention task with fully predictable stimulus timing (**Figure 1**). Observers (n = 10 × 2 sessions) directed temporal attention to one of two sequential grating targets separated by a 300 ms stimulus onset asynchrony (SOA) and discriminated the orientation of one grating on each trial. A precue tone before the targets instructed observers to attend to either the first target (T1) or second target (T2), and a response cue tone after the targets instructed observers to report the tilt (clockwise or counterclockwise) of one of the targets. The response cue matched the precue with 75% validity, so observers had incentive to direct their attention to different points in time according to the precue.

**Figure 1.**
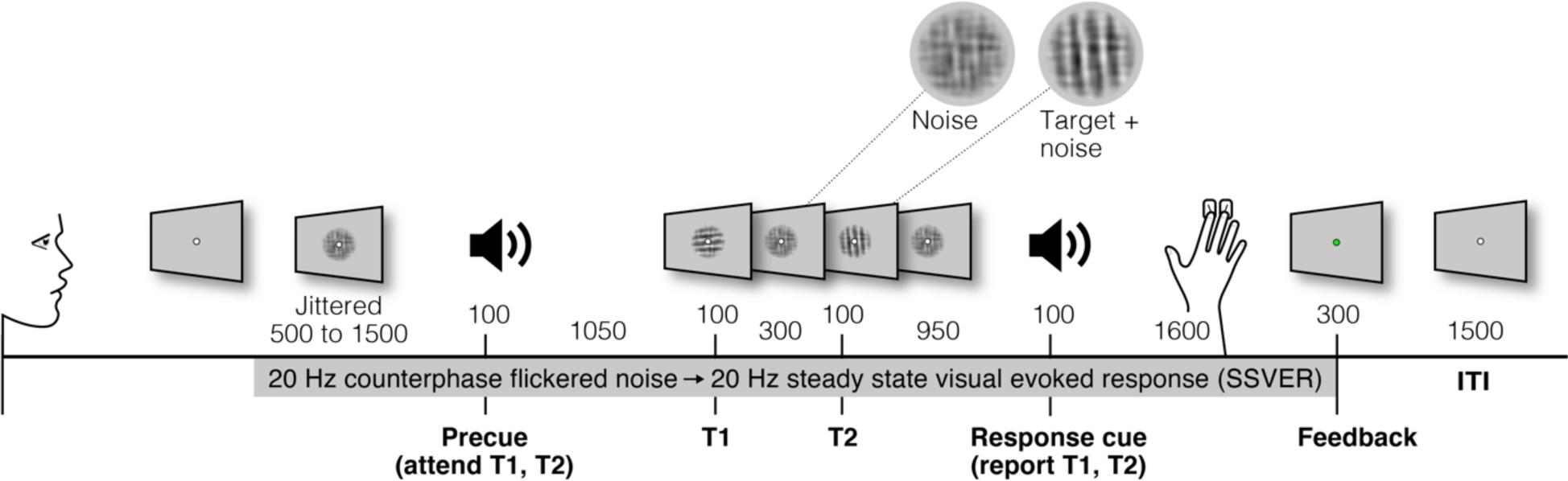
Temporal cueing task. Trial timeline showing stimulus durations and SOAs in ms. Targets were embedded in 20 Hz counterphase flickering noise. Precues and response cues were pure tones (high = cue T1, low = cue T2).

Temporal attention improved orientation discrimination performance, consistent with previous findings (3, 4, 13, 30, 56). Perceptual sensitivity (*d’*) was higher overall for valid trials (when the response cue matched the precue) than invalid trials (when the response cue did not match) (main effect of validity: F(1,9) = 8.03, *p* = 0.02, 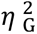 = 0.25) (**Figure 2**). Perceptual sensitivity for T1 was higher than that for T2 (main effect of target: F(1,9) = 16.47, p = 0.0028, 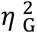 = 0.26). No other main effects or interactions were found for *d’* (F(1,9) < 3.05, p > 0.11). Given the directional nature of time, we also tested how temporal attention affected each target separately. The improvement in *d’* with temporal attention was significant for T1 individually (F(1,9) = 9.08, *p* = 0.01, 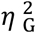 = 0.38; valid: mean *d’* = 1.79, SD = 0.44; invalid: mean *d’* = 1.21, SD = 0.88) with a trend for improvement for T2 (F(1,9) = 4.75, *p* = 0.06, 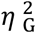 = 0.20; valid: mean *d’* = 1.20, SD = 0.39; invalid: mean *d’* = 0.85, SD = 0.61). Temporal attention did not affect observers’ discrimination criterion, or bias to report clockwise vs. counterclockwise (F(1,9) = 0.67, p = 0.43). Reaction time (RT) was faster for valid versus invalid trials overall (F(1,9) = 39.87, *p* < 0.001, 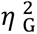 = 0.76) and for each target individually ([T1: F(1,9) = 43.02, *p* < 0.001, 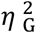 = 0.79; valid: mean RT = 0.62 s, SD = 0.23 s; invalid: mean RT = 0.89 s, SD = 0.23 s], [T2: F(1,9) = 34.71, *p* < 0.001, 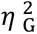 = 0.74; valid: mean RT = 0.63 s, SD = 0.24 s; invalid: mean RT = 0.88 s, SD = 0.22 s]). No other main effects or interactions were found for RT (F(1,9) < 2.66, p > 0.14) (**Figure 2**). Thus, we can rule out any speed-accuracy tradeoffs in the effect of the precue on performance.

**Figure 2.**
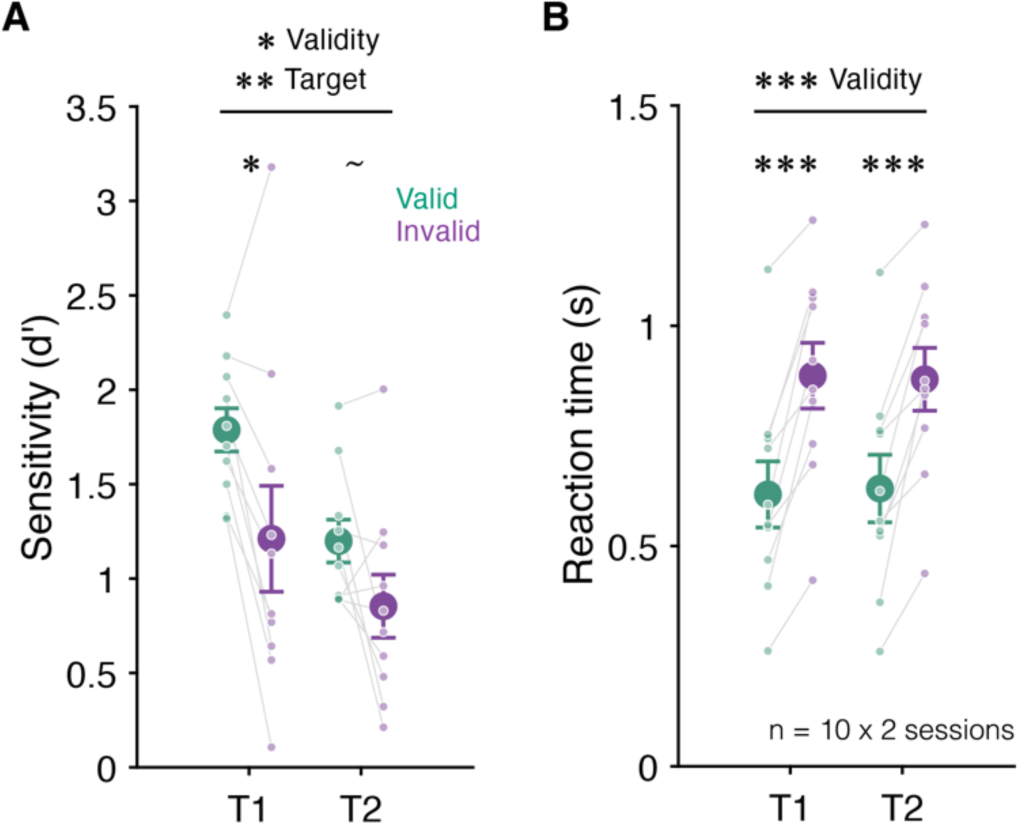
Behavior. Mean (**A**) perceptual sensitivity and (**B**) reaction time for each target (T1, T2) by attention condition. Sensitivity was higher and reaction time faster for valid (green) than invalid (purple) trials. Error bars indicate ± 1 SEM. ∼ p < 0.1; * p < 0.05; ** p < 0.01; *** p < 0.001.

### Continuous tracking of visual responses using ITPC

To allow continuous tracking of visual responses as observers deployed temporal attention, we used time-resolved SSVER. The grating targets were embedded in a stream of 20 Hz counterphase flickering noise, filtered to have similar orientation and spatial frequency content as the targets. The targets were presented superimposed on the noise background for 100 ms each and also flickered at 20 Hz in phase with the ongoing noise flicker. This protocol allowed visual cortical responses to be tracked throughout the trial by measuring the 20 Hz SSVER. The SSVER technique allows stimulus-driven activity to be isolated and measured with high signal-to-noise (57). The use of a high-frequency flicker allows high temporal-resolution measurements of the SSVER signal and increases the localization of the signal to posterior sites (48, 58).

For each observer, we first measured the 20 Hz SSVER power for each channel, calculated from the average time series across all trials (henceforth “trial-average power”), and selected the five most visually responsive channels for further analysis. As expected, the most visually responsive channels were located in the back of the head, consistent with occipital responses (**Figure 3A**). We confirmed strong 20 Hz trial-average power in these channels and found that power was enhanced specifically at the 20 Hz stimulus frequency (**Figure 3C**).

**Figure 3.**
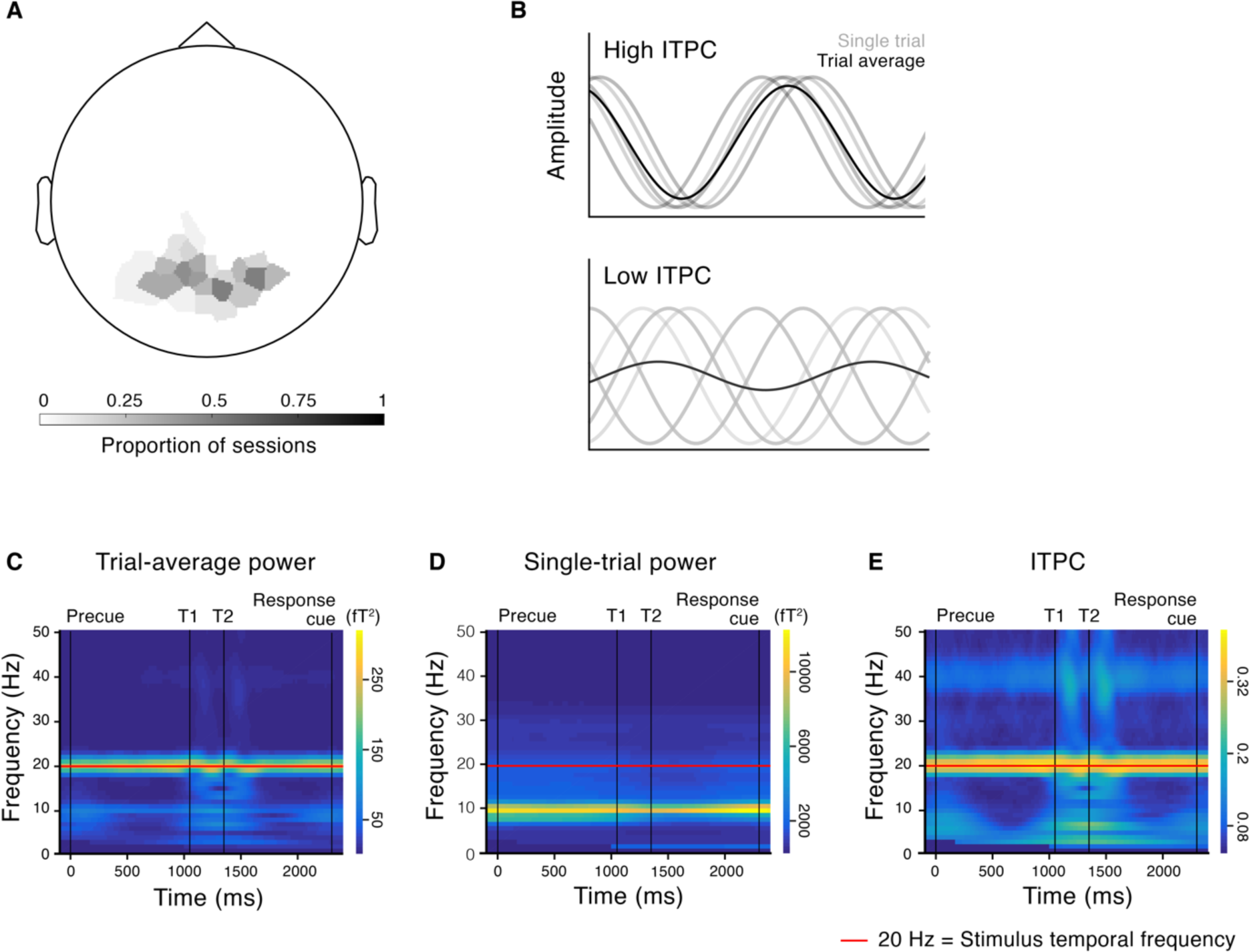
MEG responses to steady-state visual stimulation. (**A**) Topography of visually responsive channels across observers (top five channels selected for each session based on 20 Hz trial-average SSVER power). Grayscale indicates the proportion of sessions across observers in which the channel was selected as visually responsive. (**B**) Illustration of cartoon time series from five trials (gray) and their average (black). High phase coherence across trials results in high ITPC and trial-average power (top); low phase coherence across trials results in low ITPC and trial-average power (bottom), despite the same single-trial power. (**C-E**) Time frequency spectrograms of (**C**) trial-average power reveals a strong 20 Hz signal at the stimulus frequency. Trial-average power depends on two factors: (**D**) the power on single trials and (**E**) phase coherence across trials, or ITPC. The trial-average power at 20 Hz was more strongly reflected in ITPC than in single-trial power.

Trial-average power depends on two factors: 1) single-trial power and 2) phase coherence across trials at the frequency of interest, or ITPC (**Figure 3B**). Therefore, we next asked how the 20 Hz SSVER depended on these two factors. As in previous studies that have pulled apart ITPC and single-trial power (51–54), we found that ITPC was the dominant component of the trial-average SSVER, with a strong and specific 20 Hz ITPC response (**Figure 3E**). Single-trial power had a relatively weak 20 Hz component and was dominated by alpha power around 10 Hz (**Figure 3D**). We therefore used ITPC as our primary measure to quantify responsiveness to visual stimulation in our subsequent analyses.

### Temporal expectation was accompanied by a ramping increase in ITPC

We first asked how temporal expectation affected ITPC in the period between the precue and the first target. In all trials, a predictable interval of 1.05 s elapsed between the onset of the precue and the onset of T1, so observers could form an expectation about the timing of T1 onset regardless of which target was precued to be attended (4, 13). We observed an apparent neural signature of expectation in the form of a slow ramping increase of ITPC (mean slope = 0.047 ΔITPC/s, SD = 0.043) during the precue-to-T1 period (**Figure 4A, Supplementary Figure 1**). To quantify the rate of increase, we fit lines to each observer’s ITPC time series from the onset of the precue to 80 ms before T1 (a cushion before T1 was inserted to prevent responses to T1 from interfering with the fit, see Methods, Model fitting and parameter estimation). The slope of the ITPC time series was positive for 8/10 observers and significantly greater than zero across the group (F(1,9) = 12.02, *p* = 0.007, 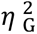 = 0.40) (**Figure 4B**). In contrast, single-trial power showed no ramping increase in the precue-to-T1 period (**Supplementary Figure 1C**), confirming that a ramping increase in cortical response reliability, as opposed to improved phase estimates due to increased signal strength, was the critical factor driving the change in ITPC.

**Figure 4.**
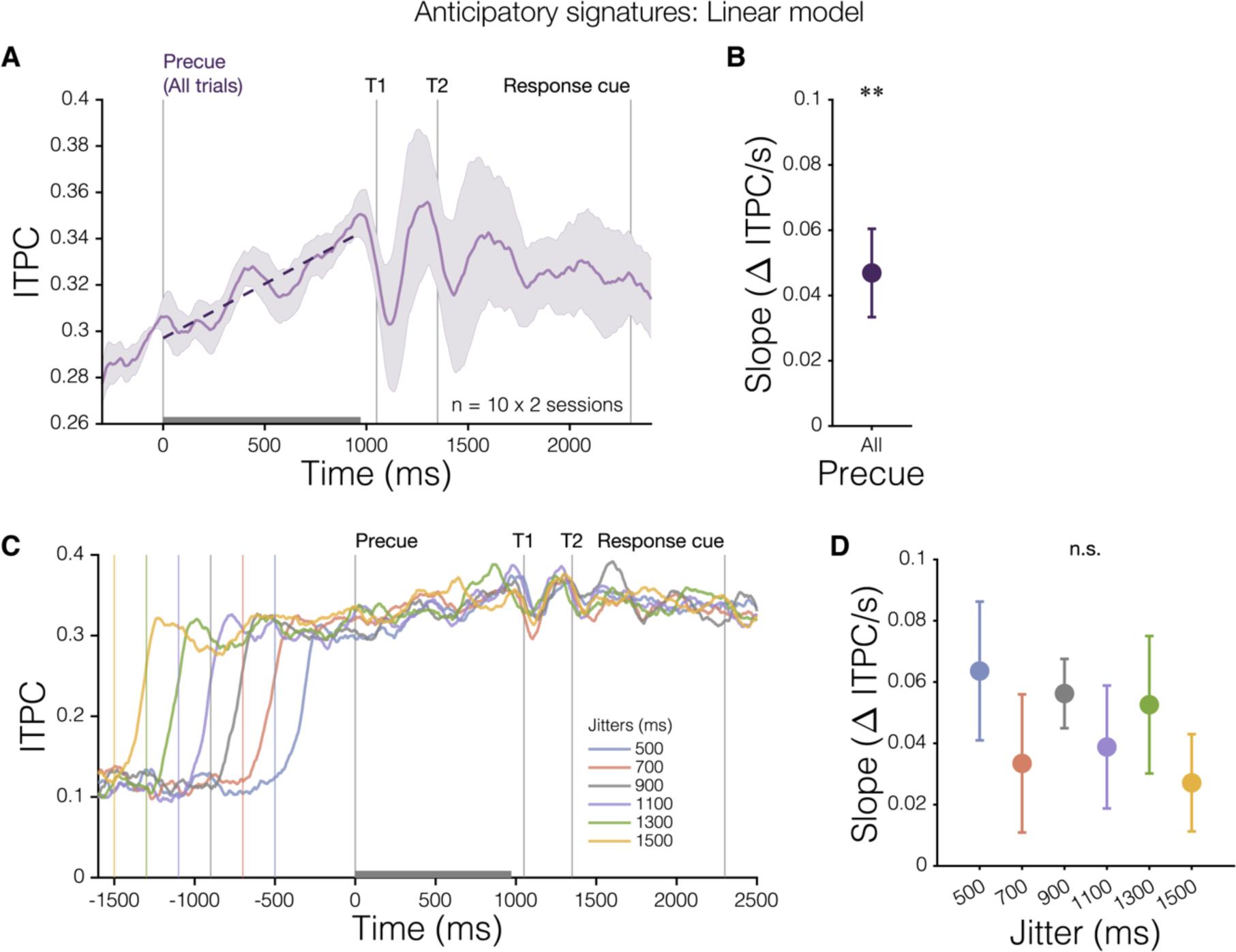
Anticipatory ITPC ramping associated with temporal expectation. (**A**) Group 20 Hz ITPC time series across all trials (*n* = 10). Lines were fit to each observer’s ITPC time series during the precue-to-T1 period (gray bar). Linear fit to group data (dashed purple line) shown for visualization. (**B**) Mean fitted slopes were significantly greater than zero. (**C**) ITPC ramp is time-locked to the precue. Mean 20 Hz ITPC time series (*n* = 10) for each jittered duration of SSVER stimulation before the precue. Colored vertical lines indicate the onset of the stimulus flicker for each duration. (**D**) ITPC slopes during the precue-to-T1 period did not differ across jitter durations. Error bars indicate ±1 SEM. ** p < 0.01 compared to zero.

To ensure this ITPC ramp was locked to the precue and not simply the continuous build-up of the SSVER signal, we separated trials according to the duration of the SSVER stimulation period before the precue, which had been jittered from 500 to 1500 ms to prevent predictable precue timing. A one-way ANOVA revealed no significant differences in ITPC slope across the six jitter durations (F(1,9) = 1.19, p = 0.30, 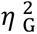 = 0.03). (**Figure 4C,D**). Therefore, the ITPC ramp was evidently initiated by the precue and indicates a gradual increase in visual cortical response reliability tied to temporal expectation.

### Delta modulation of anticipatory ITPC

Given previous findings of both linear ramping and low-frequency modulations related to temporal expectation (41, 59), we next investigated whether the pre-target ITPC time series contained any periodic activity above what is expected from the intrinsic scale-free properties of neural activity, which follows a 1/f power law (60, 61). We found that the power spectrum of the ITPC time series exceeded the 1/f expectation for a range of low frequencies, from 1 to 6 Hz (delta and low-theta range) (**Figure 5A**) (see Methods, FFT). Fitting a linear plus periodic (sine wave) model in which frequency was a free parameter showed that frequencies between 0.5-3 Hz explained the most variance in the periodic component of the ITPC time series for all observers and precue conditions, with a mean of 1.69 Hz for precue condition fits to individual data and 2.13 Hz for the group fit to all trials (**Figure 5B,C**). Thus, visual response reliability during the predictable pre-target interval exhibited periodic modulation, primarily at delta frequencies, on top of the ramping modulation.

**Figure 5.**
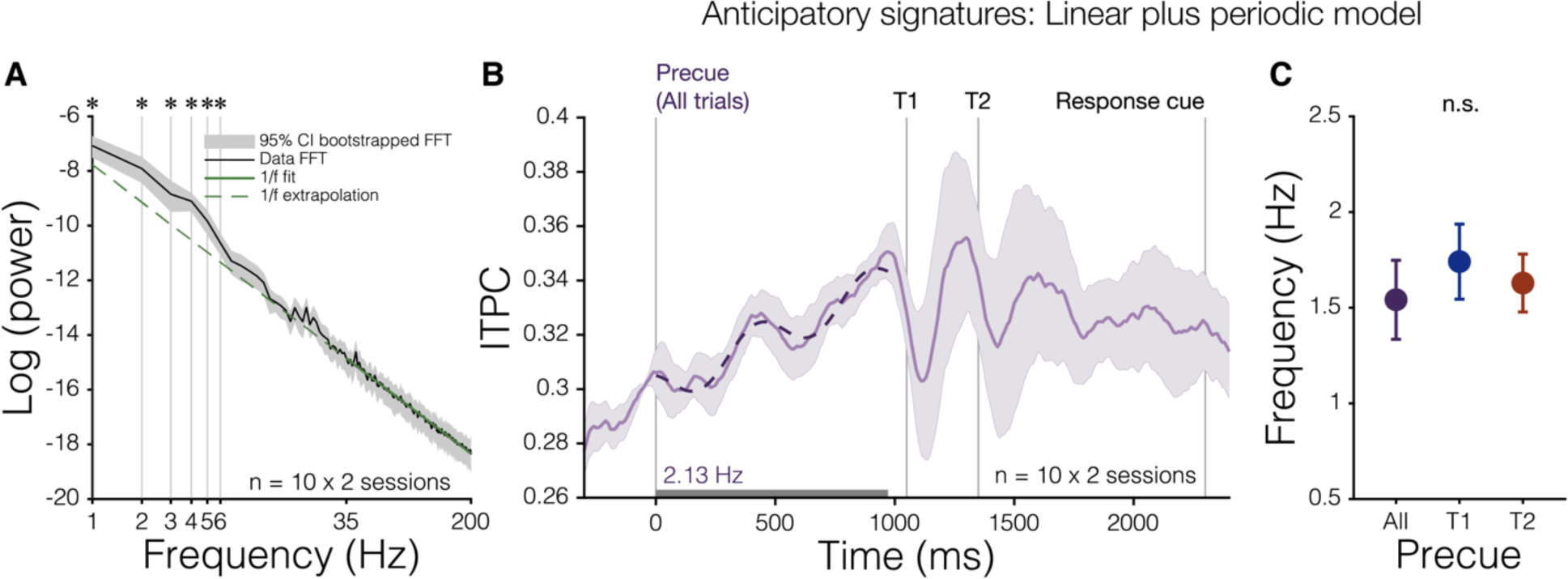
Anticipatory periodic modulations of ITPC. (**A**) The predictable precue-to-T1 period (gray bar in B) contains low frequency (1-6 Hz) periodicity above that expected from 1/f scale-free activity, determined by a linear fit to the power from 35-200 Hz (solid green line) in log-log space and extrapolated to lower frequencies (dashed green line). (**B**) Group 20 Hz ITPC time series from all trials fit with a linear plus periodic model (dashed line), in which frequency was a free parameter, indicates a dominant modulatory frequency near 2 Hz. Shaded error bars show ±1 SEM, normalized by a 100 ms baseline period before the precue. (**C**) Fitted frequencies to individual observer 20 Hz ITPC time series were similar across precue conditions. Error bars indicate ±1 SEM.

### Temporal attention phase-shifts the periodic modulation of anticipatory ITPC

Next, we tested the effect of temporal attention on the ramping and periodic components of the pre-target ITPC time series. To do so, we fit linear plus periodic models to each precue condition and assessed differences in fitted parameters. To determine whether temporal attention impacted the phase of the periodic modulation, we fixed frequency across observers and conditions and tested each of the six frequencies (1-6 Hz) that had exceeded the 1/f expectation (**Figure 5A**). The 2 Hz model fit (**Figure 6**) showed a significant phase difference between precue conditions, with a mean phase shift of 157° (*κ* = 0.43, Rayleigh z = 5.13, p = 0.0035 uncorrected, p = 0.021 after Bonferroni correction for multiple comparisons across 6 frequencies) (**Figure 6C**). The 2 Hz model also showed significant phase clustering in precue T1 trials (mean phase = 183°, *κ* = 1.83, Rayleigh z = 4.72, p = 0.0059 uncorrected, p = 0.035 after Bonferroni correction). The amplitude of the 2 Hz component and the slope and intercept of the linear component were all unaffected by the precue condition (all p > 0.1 uncorrected) (**Figure 6B**). The phase shift was specific to 2 Hz, which concords with the dominant frequency observed in the time series (**Figure 6A**); none of the other models using different fixed frequencies showed significant phase shifts (all p > 0.05 after Bonferroni correction). Interestingly, the 157° phase shift of a 2 Hz signal corresponds to a time interval 218 ms, which is similar to the SOA between the two targets. It is also similar to the SOAs at which maximal attentional tradeoffs occur: in a voluntary temporal attention study (Denison et al., 2021) that varied the interval between the two targets, the attentional precue had the largest effect on perceptual sensitivity when the targets were separated by 250 ms. Taken together, it appears that temporal expectation elicits a periodic modulation of visual response reliability in the delta band, and temporal attention shifts the phase of this modulation depending on the precise timing of the task-relevant target.

**Figure 6.**
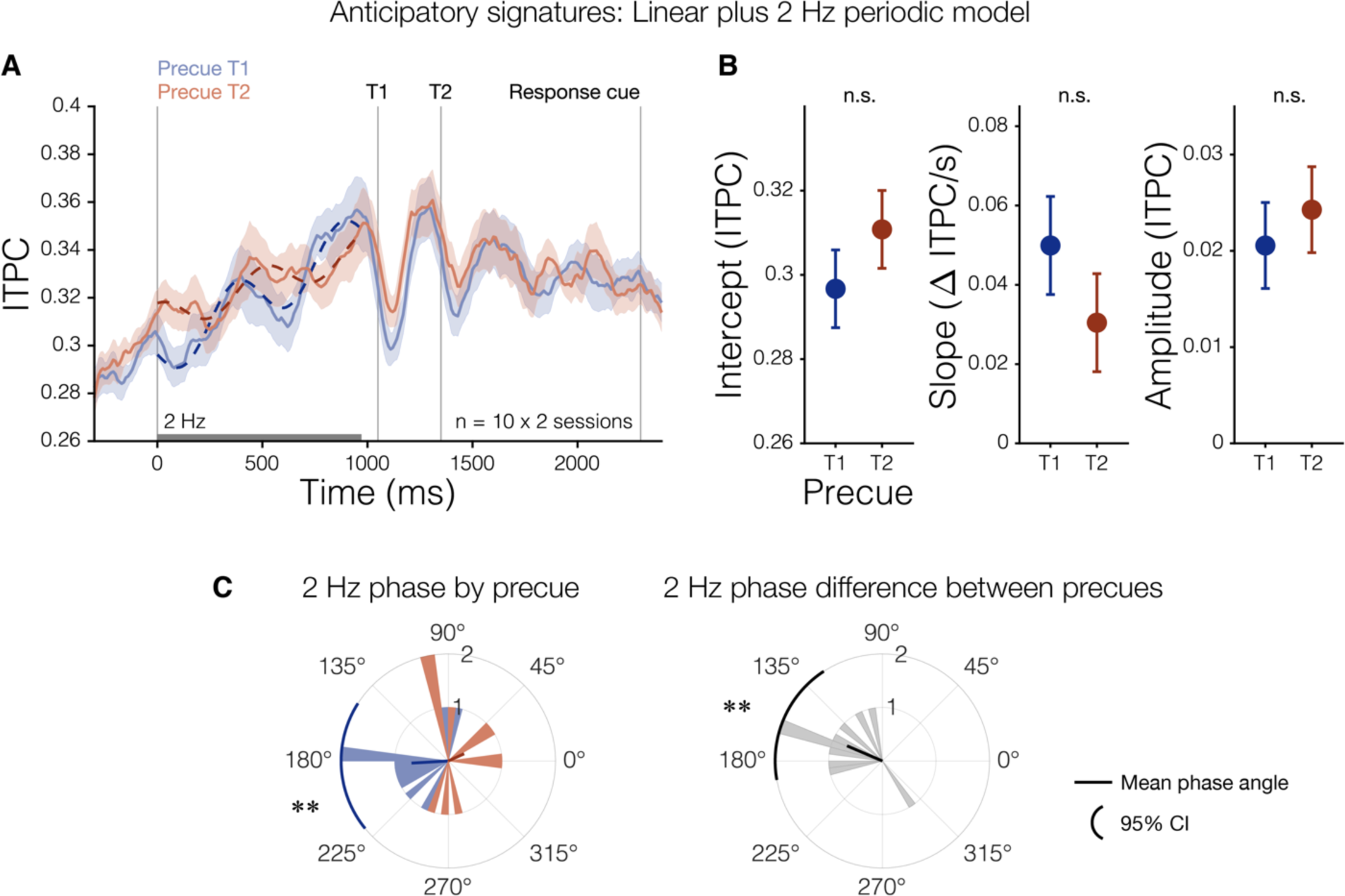
Temporal attention phase-shifts anticipatory ITPC modulation. (**A**) Group 20 Hz ITPC time series for each precue condition, fit with linear plus 2 Hz periodic models (dashed lines) to the precue-to-T1 period (gray bar). Shaded error bars indicate ±1 standard error of the difference (SED). (**B**) No difference in the fitted intercept, slope, or amplitude between precue conditions. Error bars indicate ±1 SED. (**C**) The 2 Hz phase is significantly concentrated on precue T1 trials (left), and temporal attention phase-shifts the periodic modulation (right). Shaded histograms show phase across observers. ** p < 0.01 uncorrected, p < 0.05 after Bonferroni correction.

### Temporal attention transiently affected the target-evoked ITPC response

We next asked how temporal attention affected the ITPC response to the targets. A target stimulus briefly superimposed on an SSVER stimulus can either enhance or suppress the ongoing SSVER, such that some observers show transient increases (upward peaks) in the SSVER signal following the target stimulus whereas others show transient decreases (downward peaks) (**Supplementary Text**). Thus, we asked how temporal attention affected the magnitude of this target-evoked ITPC response for each observer using a normalized measure of ITPC (see Methods, Peak analysis). The group average normalized ITPC time series has clear peaks after each target (**Figure 7A**).

**Figure 7.**
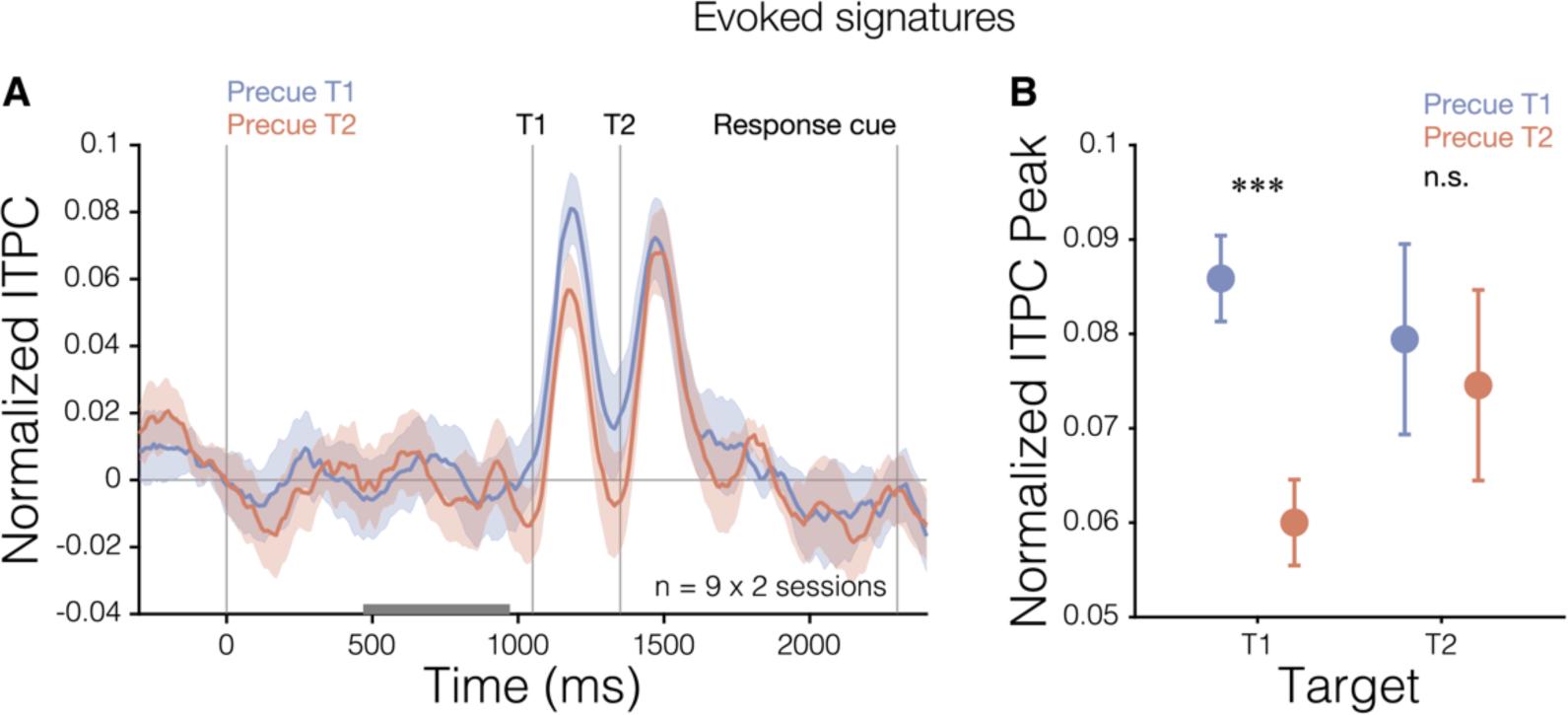
Group normalized ITPC. (**A**) Group normalized 20 Hz ITPC time series for each precue condition. Gray bar indicates the 500 ms baseline period used for normalization. (**B**) Effect of temporal attention on normalized target-evoked ITPC responses measured at individual observer peak times. Temporal attention increased the target-evoked ITPC response for T1. Error bars indicate ±1 SED. *** p < 0.001.

The magnitude of the target-evoked ITPC response to T1 was larger when T1 was precued than when T2 was precued (**Figure 7A**). To quantify the magnitude of this difference, we measured the peak of the normalized target-evoked ITPC response for each individual observer for the two precue types (see Methods, Peak analysis). We found that temporal attention increased the target-evoked ITPC response to T1 (F(1,8) = 32.34, p = 0.0004, 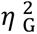 = 0.23; precue T1: mean = 0.086, SD = 0.088; precue T2: mean = 0.060, SD = 0.083 (**Figure 7B**). We found no significant effect of the precue on the response to T2 (F(1,8) = 0.23, p = 0.64, 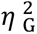 = 0.005; precue T1: mean = 0.079, SD = 0.086; precue T2: mean = 0.075, SD = 0.088). The presence of an effect of the precue for T1 and absence for T2 was reflected in a significant interaction of target and validity (F(1,8) = 7.17, p = 0.028, 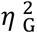 = 0.055). See **Supplementary Table 1** for all ANOVA results for the target-evoked response analyses.

We further investigated the effect of the precue on other measures of the evoked response to the targets. At the same peak times, trial-average normalized SSVER power behaved similarly to ITPC, whereas single-trial normalized 20 Hz power was unrelated to precue validity (**Supplementary Table 1**). Specifically, temporal attention increased the trial-average normalized SSVER power in response to T1 (F(1,8) = 5.80, p = 0.043, 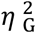 = 0.11; precue T1: mean = 560 fT^2^, SD = 560 fT^2^; precue T2 mean = 407 fT^2^, SD = 454 fT^2^) but not T2 (F(1,8) = 0.22, p = 0.65, 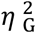 = 0.0026; precue T1: mean = 406 fT^2^, SD = 487 fT^2^; precue T2 mean = 377 fT^2^, SD = 497 fT^2^, **Supplementary Figure 1H**). Single-trial normalized 20 Hz power, in contrast, showed no effect of precue for either target (F(1,8) < 2.05, p > 0.19, **Supplementary Figure 1I**), demonstrating that the change in ITPC was not driven by a change in 20 Hz power regardless of phase. Finally, we confirmed that the ITPC effect for T1 was specific to 20 Hz (**Supplementary Figure 2**). Therefore, directing temporal attention to the first target changed the steady-state response to that target in a transient fashion.

## Discussion

We investigated how temporal attention dynamically modulates visual processing by using time-resolved SSVER to continuously track visual cortical activity as observers attended to precise points in time. Our two-target temporal cueing task (3, 4, 10, 13) allowed us to measure specific effects of voluntary temporal attention to task-relevant time points, over and above the effects of temporal expectation due to predictable stimulus timing. Temporal attention affected cortical response reliability both in anticipation of and in response to target stimuli. Visual response reliability slowly ramped up as target onset approached, an overall effect of temporal expectation. In addition, response reliability exhibited low-frequency modulations before target onset, which shifted in phase according to which target was attended. Temporal attention to task-relevant time points also enhanced target-evoked responses, specifically for the first target. These findings demonstrate both anticipatory and evoked visual cortical signatures of voluntary temporal attention, when temporal expectation is controlled.

Both ramping activity and neural oscillations have been proposed to serve as timing mechanisms in the brain (59, 62), and both have been observed in anticipation of a predictable stimulus. Studies of temporal expectation have found anticipatory ramps in EEG voltage known as the contingent negative variation (CNV), often when a speeded response to a target was required (28, 63–67). The CNV is usually associated with response preparation or interval timing (68) and has been localized to frontal and parietal areas (69). Ramping activity in motor and parietal areas has also been observed leading up to timed movements (35, 70). In visual cortex, gamma responses to an irrelevant grating ramped up over the course of ∼1 s before the expected target time, and alpha power decreased (42).

Temporal expectation has also been associated with low-frequency oscillations in the delta band (∼0.5-4 Hz) in both sensory (38, 41, 71, 72) and fronto-parietal (73, 74) areas. A study in macaques (38) found that interleaved rhythmic auditory and visual stimuli elicited V1 delta modulation, which phase-shifted to align with input in the task-relevant modality. Delta phase was related to the amplitude of visual responses trial by trial, suggesting that delta can modulate response gain. Alignment of rhythmic stimuli to a preferred phase of delta also has behavioral consequences, improving discrimination (40, 72, 74) and speeding reaction times (38, 39, 74). These reports of delta activity come from experiments that also presented rhythmic stimuli at delta frequencies. Thus, it is possible that the activity was due to stimulus-driven entrainment (38, 40, 71). However, other expectation experiments found delta phase synchronization even in the absence of rhythmic input, using predictable cue interval timings (41, 74). None of the above-mentioned studies attempted to isolate the effects of voluntary temporal attention from those of temporal expectation.

Here, we found expectation-related ramping and low-frequency modulation in the reliability of stimulus-specific visual responses, which was phase-shifted by temporal attention. The time-resolved SSVER approach revealed a direct effect of ramping and periodic modulations on cortical responses to visual stimuli presented at each time point, demonstrating the continuous functional impact of these modulatory signals during target anticipation. Given that the ITPC signal and attention-related modulations were narrow-band at the stimulus frequency of 20 Hz and phase-locked to the stimulus, these measurements very likely reflect neural responses driven by the stimulus flicker rather than endogenously generated activity in the beta band. Unlike in previous studies (26, 27, 42) no speeded responses to targets were required, ruling out the possibility that the cortical modulations were associated with response preparation (64). SSVER studies of spatial attention sustained across seconds have reported SSVER amplitudes that are relatively constant across a trial (51, 52), suggesting that the ramping plus periodic pattern we observed is specific to situations when the observer can predict the upcoming target time. Interestingly, an overall anticipatory effect of temporal expectation that is time-shifted by temporal attention is similar to the pattern we have observed for microsaccades, which also provide continuous measures of anticipatory processes (10, 14).

We also found that temporal attention enhanced the evoked ITPC response even for fully predictable stimuli, but this enhancement occurred only for T1 and not for T2. One possible explanation, albeit speculative, is that the neural mechanisms of temporal attention differ for T1 and T2 given the directionality of time. When stimuli appear in rapid succession, anticipatory signals that slowly build up over time would mainly affect the first stimulus in the sequence. It may be that at occipital stages, changing the relative responses to the two targets by modulating T1 alone is sufficient to bias downstream competition between the two target representations and produce attentional tradeoffs in behavior. In our data, consistent with a relative modulation of target responses by attention, the T2 evoked ITPC response was higher than the T1 response when T2 was precued and lower than the T1 response when T1 was precued.

An investigation of temporal expectation using SSVER yielded mixed results (50): post-target SSVER amplitude was higher for predictable vs. unpredictable stimuli for some flicker frequencies but lower for other flicker frequencies, with no increase in SSVER amplitude in advance of a predictable target time. However, the SSVER stimulus had no spatial overlap with the target location, so any expectation-related modulation specific to the target location would not have been measurable in the SSVER signal. Here, with spatially co-localized SSVER and target stimuli, temporal attention increased the impact of the targets on the SSVER for all observers. However, the targets themselves increased the SSVER for some observers and decreased it for others. This directionality was stable across sessions for each observer. To our knowledge this individual variation is an unexplained observation in the field that deserves further study (**Supplementary Text**).

Our normalization model of dynamic attention predicts that temporal attention will modulate sensory responses (3). Based on fits to psychophysical data, the model also predicts that the effects of temporal attention on neural gain will be transient and roughly coincident with the sensory responses. The current data are consistent with both model predictions: sensory modulation by temporal attention and modulation at a transient timescale. Specifically, the model predicts a change in gain at the neuronal level, which requires a linking hypothesis to make predictions for MEG measures. Higher gain would increase the probability or rate of neuronal spiking in response to each stimulus flicker, which in turn would be expected to lead to more reliable population responses to the SSVER stimulus across trials and thus higher ITPC. Such a mechanism also might be expected to increase single-trial power, which we did not observe. However, the SSVER stimulus itself did not substantially increase 20 Hz single-trial power above baseline. This raises the possibility that stimulus-driven 20 Hz activity interacted with ongoing endogenous (non-phase locked) 20 Hz power, which may have obscured gain changes in the single-trial power measure. Alternatively, changes in ITPC could have arisen from a mechanism other than gain changes, such as more consistent timing of stimulus-driven responses without changes in neuronal response amplitude. Invasive recordings, and perhaps modeling efforts, could distinguish between these possibilities.

This linking hypothesis between gain increases at the neuronal level and ITPC increases in the SSVER can account for the pre-target effects of attention and expectation we observed, as well as post-target effects for those observers showing a transient increase of the ITPC following each target. However, transient reductions of the ITPC in response to target stimuli, regardless of attention (**Supplementary Text**), appear to arise from observer-specific interactions between the ongoing SSVER and target-evoked responses and are thus not predictable from the normalization model of dynamic attention (3), which does not include specific biophysical structure. The model also could not predict the anticipatory ramping and periodic modulation we observed here, because the behavioral data alone could not distinguish anticipatory from stimulus and post-stimulus processing; future versions of the model should incorporate these new experimental findings.

## Conclusions

Using time-resolved SSVER, we studied how voluntary attention to points in time continuously impacts the reliability of cortical responses to visual stimulation. Our measurements revealed two complementary neural mechanisms of temporal attention: (1) a slow, periodic fluctuation of visual cortical ITPC that anticipated predictable target times and shifted in time depending on which time point was to be attended, and (2) a transient, post-target increase of the evoked ITPC response to targets at attended time points, which was specific to the first target. This transient modulation changed the relative strengths of the two target responses as a function of attention. By revealing the full timecourse of temporal attentional modulations of sensory activity, the results show how attention dynamically alters visual signals to select and prioritize visual information at precise moments in time.

## Methods

### Observers

Ten observers (5 females), ages 24-35 years (M = 28.5 years, SD = 4.2 years) participated, including authors RND and KJT. Each observer completed two MEG sessions, which allowed us to check for consistency across sessions for each observer while collecting 20 sessions of MEG data in total, comparable to previous MEG studies of vision (75, 76). The sample size was also similar to previous temporal attention studies that measured accuracy (4, 13, 29) and to several studies on spatial attention (77–79). All observers provided informed consent, and the University Committee on Activities involving Human Subjects at New York University approved the experimental protocols. All observers had normal or corrected-to-normal vision.

### Stimuli

Stimuli were generated on an Apple iMac using MATLAB and Psychophysics Toolbox (80, 81) and were displayed using a gamma-corrected InFocus LP850 projector (Texas Instruments, Warren, NJ) with a resolution of 1024 × 768 pixels and a refresh rate of 60 Hz. Stimuli were projected via a mirror onto a translucent screen at a viewing distance of 42 cm. Stimulus timing accuracy was confirmed with photodiode measurements. Stimuli were presented on a medium gray background (206 cd/m^2^). A central fixation circle subtended 0.15° visual angle.

#### SSVER stimulus

Throughout the trial, a 20 Hz contrast-reversing flickering noise patch was presented foveally, with the fixation circle overlaid. The noise patch was 4° in diameter with a blurred outer edge subtending 0.4°, fading to the background gray. To create the noise patch, pixel noise was generated from a normal distribution then bandpass filtered in both orientation and spatial frequency around target values: in orientation 0° ± 5° and 90° ± 5° forming a vertical and horizontal crosshatch pattern, and in spatial frequency 1.5 cpd ± 1 octave. A new noise patch was randomly generated on each trial and then flickered throughout the trial with a duty cycle of 33% (1 frame of the original noise image in alternation with 2 frames of the contrast-reversed image). The contrast of the SSVER stimulus was adjusted to ensure high visibility of the target stimuli while still producing a clear SSVER signal (40-55% noise patch contrast across observers).

#### Targets

Visual targets were 1.5 cpd sinusoidal gratings of 4° in diameter with a blurred outer edge subtending 0.4°. Targets were also presented foveally and were added pixelwise to the noise patch stimulus. To match the temporal frequency of the targets to the SSVER stimulus, the targets were also contrast-reversed for 2 cycles at 20 Hz, in phase with the noise patch flicker, for a total target duration of 100 ms. Target grating contrast was adjusted so that, when added to the noise patch, the total contrast was 100% (grating contrast 45-60% across observers).

#### Cues

Auditory cues were pure sine wave tones 100 ms in duration (with 10 ms cosine amplitude modulation ramps at onset and offset) presented through earbuds at a comfortable volume. There were two possible auditory cue tones, one high-pitched (1046.5 Hz, C6) and one low-pitched (440 Hz, A4).

### Experimental procedure

All observers participated in two 2-hour MEG sessions on separate days, on average 3.6 days apart. Each session included 12 experimental blocks approximately 6 minutes each in duration. Observers took breaks between blocks and pressed a button to indicate their readiness for the next block.

#### Task

On each trial, observers were asked to discriminate the orientation of one of two grating targets (**Figure 1**). The two targets (T1 and T2) were each presented for 100 ms, separated by a stimulus onset asynchrony (SOA) of 300 ms, based on previous psychophysical studies (3, 4, 13). Each target was tilted slightly clockwise (CW) or counterclockwise (CCW) from either the vertical or horizontal axis, with independent tilts and axes for T1 and T2. Both vertical and horizontal axes were used to reduce the number of trials on which the two stimuli were identical and discourage observers from adopting a strategy of judging whether the stimuli were the same or different to aid in discrimination. An auditory precue 1,050 ms before the first target instructed observers to attend to the first or the second target (high tone: attend T1; low tone: attend T2). An auditory response cue 950 ms after T2 indicated which of the two targets’ tilt to report (high tone: report T1; low tone: report T2). Observers pressed one of two buttons to indicate whether the tilt was CW or CCW relative to the main axis within a 1600 ms response window. The response cue matched the precue on 75% of the trials (valid trials) and mismatched the precue on the remaining 25% (invalid trials). At the end of each trial, feedback was provided by a change in color of the fixation circle (green: correct; red: incorrect; blue: response timeout). At the end of each block, performance accuracy (percent correct) was displayed. The timing of auditory and visual events was the same on every trial. From trial to trial, the allocation of temporal attention varied (depending on the precue), and the response selection varied (depending on the response cue).

So that only the precue would provide predictive timing information about the target onsets, and to allow time for the SSVER signal to stabilize before any trial events occurred, the interval between the SSVER onset and the precue was jittered from 500 to 1500 ms in steps of 200 ms. Between each trial, an inter-trial interval of 1500 ms was presented, during which only the fixation circle appeared on a gray background. Observers were encouraged to blink during this period. The fixation circle turned black at the start of the ITI, turned white 500 ms before the SSVER onset to indicate that the trial was about to begin, and remained white throughout the trial until the response feedback was provided.

Each session consisted of 12 blocks of 41 trials each with all combinations of cue type (valid: 75%, invalid: 25%), probed target (T1, T2), target tilt (CW, CCW; independent for T1 and T2), and target axis (horizontal, vertical; independent for T1 and T2) in a randomly shuffled order. To reduce adaptation to the SSVER stimulus, every fifth trial was a blank trial, with an additional blank trial at the beginning of each block. This procedure yielded a total of 768 temporal attention trials per observer across the two sessions.

#### Training and thresholding

Prior to MEG, observers completed at least one session of training to familiarize them with the task and to determine their orientation tilt thresholds. During the training session, a chin-and-head rest was used to hold head position and viewing distance constant. Stimuli were displayed on a gamma-corrected Sony Trinitron G520 CRT monitor with a refresh rate of 60 Hz at a viewing distance of 57 cm. A 3-up-1-down staircase procedure was used to estimate each observer’s tilt threshold to achieve ∼79% accuracy across all trials. Threshold tilts ranged from 1.5-5°.

#### Eyetracking

Eye position was recorded using an EyeLink 1000 eye tracker (SR Research) with a sampling rate of 1000 Hz. Raw gaze positions were converted into degrees of visual angle using the five-point-grid calibration, which was performed at the start of each MEG session.

#### MEG recording

Data were recorded continuously in each block with a 157-channel axial gradiometer system (Kanazawa Institute of Technology, Kanazawa, Japan) in the KIT/NYU facility at New York University. The magnetic fields were sampled at 1000 Hz with online DC filtering and a 200 Hz low-pass filter. Three orthogonally-oriented reference magnetometers placed approximately 20 cm away from the recording array recorded environmental noise.

Before recording, each observer’s head shape was digitized with a handheld FastSCAN laser scanner (Polhemus, VT, USA). Digital markers were placed on the forehead, nasion, and left and right tragus and peri-auricular points. To record marker locations with respect to the MEG channels, electrodes were placed on five of the locations identified by the digital markers (three points on the forehead and left and right peri-auricular points), and marker locations were measured at the beginning and end of the MEG recording session.

### Data analysis

#### Preprocessing

Data was preprocessed in MATLAB using the FieldTrip toolbox for EEG/MEG-analysis (82). Environmental noise was removed from the data by regressing signals recorded from three orthogonally oriented magnetometers against the recorded data.

MEG channels in which there was no signal or excessive noise were interpolated from neighboring channels (the number of interpolated channels per recorded session ranged from 2 to 16 (1.3-10.2%), mean = 3.9 (2.5%), SD = 3.0. Trials were manually inspected and rejected for blinks and other artifacts. The number of rejected trials per session ranged from 26 to 73 (5.3-14.8%), mean = 45.4 (9.2%), SD = 14.3.

#### Channel selection

We selected visually responsive channels as the five channels per session with the strongest 20 Hz trial-average SSVER power. Topographies confirmed selected channels were in the posterior part of the head.

#### SSVER analysis

For the SSVER analyses, trials were epoched from 1500 ms before to 5200 ms after the precue. Time-frequency representations of the data were computed using wavelet convolution with the FieldTrip toolbox. For the 20 Hz SSVER frequency, Morlet wavelets had 8 cycles (127 ms temporal resolution, 5 Hz spectral resolution). For full time-frequency spectra, frequency was sampled every 1 Hz from 1-50 Hz, time was sampled every 10 ms, wavelets were tapered with Hanning windows, and the time windows scaled with frequency according to 10 s divided by the frequency.

#### Trial-average power

Trial-average power quantifies the power of a periodic signal that is phase-locked across trials and so survives averaging across trials. This measure has been commonly used in the SSVER literature (45, 83–85). To calculate the trial-average power, the time series of all trials (or all trials in a given experimental condition) were first averaged together, giving the mean time series across trials. Time-frequency analysis was then performed on this single time series to calculate the power across time at a frequency of interest. Trial-average power depends on two factors: single-trial power and ITPC.

#### Single-trial power

Single-trial power quantifies the power of a periodic signal present on individual trials. To calculate the single-trial power, time-frequency analysis was performed on each individual trial, and these time-frequency representations were then averaged together across trials. This procedure preserves oscillatory activity that is not phase-locked across trials but relies on having sufficient signal-to-noise at the single trial level to measure the periodic signal.

#### ITPC

Inter-trial phase coherence (ITPC) quantifies the phase alignment of a periodic signal across trials. The measure ranges from 0 (no phase alignment) to 1 (perfect phase alignment) (86). To calculate ITPC, time-frequency analysis was performed on each individual trial, returning estimates of the instantaneous phase angle at each point in time for a frequency of interest. The following standard formula was then used to calculate the alignment of these phase angles across trials, providing an estimate of ITPC:

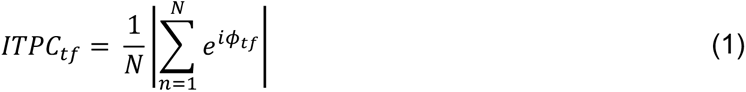

where *ϕ_tf_* is the instantaneous phase angle in radians at a time point *t* and frequency *f* for each trial *n*∊[1,…,*N*].

### Anticipatory ITPC analyses

#### FFT

To investigate whether there were any periodic modulations of the ITPC timeseries beyond the expected scale-free spectral content, we assessed whether the power spectrum of the ITPC timeseries in the precue-to-T1 period differed from a 1/f relationship. To do so, we converted the anticipatory ITPC time series per session into the frequency domain using fast Fourier transform (from 1 Hz to 200 Hz, with 1 Hz resolution). We bootstrapped the 20 resultant spectra 1000 times, resampling with replacement to generate a distribution of spectra. The 1/f power law—as frequency increases, power decays following a power-law function—simplifies to a linear relationship between power and frequency in log-log space (87). As such, we estimated the aperiodic components of the anticipatory ITPC period by fitting a line to the average empirical spectrum in log-log space from 35 to 200 Hz, while leaving out 60 Hz line noise and harmonics, following the procedure from (88), and extrapolated the fit to estimate the power expected from 1/f scale-free activity at lower frequencies (<35 Hz). We calculated the 95% confidence interval of the bootstrapped distribution of spectra using its 2.5 to 97.5 percentiles. Deviation of the confidence interval above or below the fitted line would be evidence for significantly more or less periodicity, respectively, than expected from scale-free activity alone, at an alpha level of 0.05.

Using the same bootstrapping procedure, we confirmed the spectra of the recorded time series, averaged across the top 5 visually responsive channels (see Methods, Channel selection) per session, did not contain 2 Hz power above 1/f expectations; this means the periodic 2 Hz activity we find in the precue-to-T1 period was specific to phase coherence of the SSVER.

#### Model fitting and parameter estimation

To quantify pre-target ITPC modulations for individual observers, we fit both linear and linear plus periodic models to the ITPC time series in the time window between the precue and first target. In all trials, a predictable interval of 1.05 s elapsed between the onset of the precue and the onset of T1, so observers could form an expectation about the timing of T1 onset. To prevent T1 from influencing ITPC measurements of pre-target activity due to the temporal bandwidth of the wavelet, we set a cushion before T1 based on the group average ITPC time series across all trials. The start of the cushion was the inflection point 80 ms before T1 when ITPC began to sharply decrease due to the T1 response. Thus the precue-to-T1 window was defined as the onset of the precue to 80 ms before T1 (0-970 ms).

To assess the overall effect of expectation on ITPC in this precue-to-T1 period, we fit the models on the ITPC time series calculated from all trials. To assess the effect of temporal attention, we fit the models to ITPC time series calculated from trials for each precue condition separately. In all cases, we fit the models for each observer and session and then statistically assessed the parameter estimates from these fits.

The linear plus periodic model was defined as:

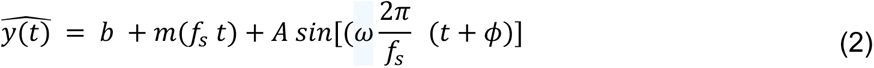

where *ŷ* is the predicted ITPC, *t* is time, *fs* is the sampling frequency (1000 Hz), *b* is the intercept, *m* is the slope of the linear component, and *A*, *ω*, and *ϕ* are the amplitude, frequency, and phase of the periodic (sine wave) component. *b*, *m*, *A*, and *ϕ* were always free parameters, and *ω* could be free or fixed to a specific frequency, e.g. 2 Hz.

The linear only model was identical but contained only the intercept and slope terms.

We estimated the model parameters by minimizing the mean squared error between the predicted and true ITPC using the optimization algorithm Bayesian Adaptive Direct Search (BADS) (89). The search bounds for each parameter were:

*b*: [0,1]

*m*: [−2, 2] ΔITPC/s

*A*: [0,1]

*ω*: [0.1, 5] Hz

*ϕ*: [−*pi*, *pi*] radians

To avoid local minima, we randomly generated 100 sets of initial parameters within the search bounds. We performed BADS optimization for each set of initial parameters, then selected the set of fitted parameters associated with the minimum error across iterations as the best fitting parameters.

#### Phase analysis

Phase analyses were performed using the Circular Statistics Toolbox (90). The effect of temporal attention on phase was assessed using Rayleigh’s z test, which tests for non-uniformity of a circular distribution.

### Evoked ITPC analysis

#### Peak analysis

To quantify the evoked ITPC response for individual observers, we estimated the latency, direction, and amplitude of the most prominent deflection in the 20 Hz ITPC following each target, which we refer to as a “peak”. As ITPC data in the selected visual channels was highly consistent across sessions for a given observer, we identified peak times for each observer using the ITPC time series averaged across the two sessions. To find the peaks, we applied the MATLAB algorithm *findpeaks.m* to the time window 0 ms to 600 ms after T1 onset, which captured the evoked ITPC responses to both T1 and T2.

All observers but one had two peaks that were either upward or downward during this window. The observer without an identifiable pair of peaks was not considered further in the peak analysis. Two observers had both upward peaks and downward peaks during the window, so for these observers we selected the pair of peaks whose timing was best aligned with the peaks of the other observers. The direction of the pair of peaks identified by this algorithm determined the classification of an observer as having either upward or downward peaks.

We next assessed the effect of attention on the magnitude of the ITPC peaks, regardless of their direction. To combine evoked ITPC data across observers for this analysis, we first normalized the data so that peaks for all observers pointed upward. To do so, we measured a baseline ITPC value for each observer and session in the window 470-970 ms after the precue (a 500 ms window before target onset, excluding the 80 ms cushion before T1, see “Slope analysis”). We then subtracted that baseline value from the time series, and for observers with downward peaks, reversed the sign:

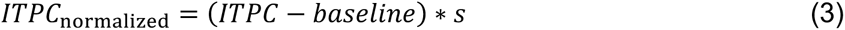

where *s* = 1 for observers with upward peaks and −1 for observers with downward peaks. This procedure allowed all evoked ITPC peaks to point upward while preserving all other features of the time series.

Finally, to quantify the ITPC magnitude at the peak times, for each observer, session, and precue condition, we took the mean normalized ITPC value in a 100 ms window centered on each peak time (i.e., following each target). Equivalent peak analyses were performed for trial-average 20 Hz power and single-trial 20 Hz power using the same peak times and normalization procedure.

#### Statistical analysis

The effects of temporal attention on behavior (*d’* and RT) and MEG time series (anticipatory model parameter estimates and evoked peak values) were assessed using repeated measures ANOVAs via the *ezANOVA* package in R. For behavioral data, the within-subject factors were session, target, and validity (valid or invalid, with respect to the match between the precue and response cue), with session as an observed factor. For MEG anticipatory analyses, within-subject factors were session and precue (precue T1 or precue T2), with session as an observed factor. For MEG peak analyses, within-subject factors were session, target, and attention (attended or unattended, with respect to the precue type, referenced to each target), with session as an observed factor. Planned ANOVAs were also conducted for each target separately.

To assess the effect of the precue on the full time series or time-frequency spectrum, we used a non-parametric test with cluster correction (91). This involved a two-step procedure using data averaged across sessions for each observer. In the first step, paired t-tests were performed at each time point or time-frequency bin (henceforth, “bin” for both) comparing precue T1 to precue T2 condition values. This operation yielded t-statistics for each bin for the empirical data. Contiguous bins passing the p < 0.05 threshold were considered to form a cluster, and a statistic was generated for each cluster by taking the sum of the t-values for bins within the cluster. The largest statistic out of all clusters was considered to be the maximum cluster statistic for the empirical data. Next, null distributions of the maximum cluster statistic were constructed by shuffling the labels for the precue conditions, computing the maximum cluster statistic for the shuffled data, and repeating this process 1000 times. Significance levels for the empirical data were determined by comparing the value of the maximum cluster statistic to the null distribution.

## Acknowledgements

We thank Sirui Liu and Luis Ramirez for assistance with early versions of the experiment. We thank Hsin-Hung Li, Amit Yashar, and members of the Carrasco and Denison Labs for helpful comments. We thank Jeffrey Walker at the NYU MEG lab for technical assistance.

## Funding

National Institutes of Health National Eye Institute R01 EY019691 (MC)

National Institutes of Health National Eye Institute F32 EY025533 (RND)

National Institutes of Health National Eye Institute T32 EY007136 (NYU)

National Defense Science and Engineering Graduate Fellowship (KJT) Boston University (RND)

## Author contributions

Conceptualization: RND, DJH, MC

Methodology: RND, KJT, DJH, MC

Investigation: RND, KJT

Visualization: RND, KJT

Supervision: RND, DJH, MC

Writing—original draft: RND, KJT

Writing—review & editing: RND, KJT, DJH, MC

## Data availability

The data that support the findings of this study are available from the corresponding author upon reasonable request.

## Supplementary materials

Supplementary Text

Tables S1 and S2

Figures S1 to S5

