## Supplementary Information for "Anticipatory and evoked visual cortical dynamics of voluntary temporal attention"

Department of Psychological and Brain Sciences
Boston University
64 Cummington Mall
Boston, MA 02215
United States

**Keywords:** temporal attention, temporal expectation, magnetoencephalography, steady state visual evoked response

#### Supplementary Tables

**Supplementary Table 1.** Peak analysis. Target-evoked 20 Hz ITPC, trial-average 20 Hz SSVER power and single-trial 20 Hz power, peak responses of normalized data

| | Statistics | | |
|---|---|---|---|
| | F(1,8) | p | $\eta^2_G$ |
| ITPC | | | |
| Session | 2.16 | 0.18 | 0.054 |
| Target | 0.24 | 0.64 | 0.004 |
| Validity | 3.89 | 0.084 ~ | 0.027 |
| Validity (T1 only) | 32.34 | 0.00046 *** | 0.23 |
| Validity (T2 only) | 0.23 | 0.64 | 0.005 |
| Session:Target | 16.66 | 0.004 ** | 0.15 |
| Session:Validity | 0.11 | 0.75 | 0.0008 |
| Target:Validity | 7.17 | 0.028 * | 0.056 |
| Session:Target:Validity | 0.22 | 0.65 | 0.005 |
| Trial-average SSVER power | | | |
| Session | 1.94 | 0.20 | 0.12 |
| Target | 1.04 | 0.34 | 0.024 |
| Validity | 2.28 | 0.17 | 0.011 |
| Validity (T1 only) | 5.80 | 0.043 * | 0.11 |
| Validity (T2 only) | 0.22 | 0.65 | 0.0026 |
| Session:Target | 22.93 | 0.0014 ** | 0.047 |
| Session:Validity | 1.39 | 0.27 | 0.002 |
| Target:Validity | 3.58 | 0.095 ~ | 0.025 |
| Session:Target:Validity | 0.14 | 0.71 | 0.0005 |

| Single-trial 20 Hz power | | | |
|---|---|---|---|
| Session | 0.80 | 0.40 | 0.009 |
| Target | 1.01 | 0.34 | 0.078 |
| Validity | 1.62 | 0.24 | 0.002 |
| Validity (T1 only) | 2.05 | 0.19 | 0.043 |
| Validity (T2 only) | 0.022 | 0.89 | 0.0003 |
| Session:Target | 3.52 | 0.098 ~ | 0.03 |
| Session:Validity | 10.35 | 0.012 * | 0.012 |
| Target:Validity | 0.74 | 0.41 | 0.003 |
| Session:Target:Validity | 1.46 | 0.26 | 0.009 |

~ $p < 0.1$, * $p < 0.05$, ** $p < 0.01$, *** $p < 0.001$

**Supplementary Table 2.** Evoked ITPC response direction

| | Direction of target-evoked ITPC response | | |
|---|---|---|---|
| | Upward, *n* = 4 | Downward, *n* = 5 | Statistics |
| Demographics | | | |
| Gender | 2 female, 2 male | 2 female, 3 male | |
| Age | Mean = 31.25 years, SD = 4.11 | Mean = 28 years, SD = 3.67 | $F(1,7) = 1.57$, $p = 0.25$, $\eta^2_G = 0.18$ |
| Handedness | all right dominant | all right dominant | |
| Behavior | | | |
| Perceptual sensitivity (*d'*) | Mean = 1.16 SD = 0.49 | Mean = 1.41 SD = 0.81 | $F(1,7) = 0.58$, $p = 0.47$, $\eta^2_G = 0.077$ |
| Reaction time (s) | Mean = 0.69 SD = 0.18 | Mean = 0.72 SD = 0.26 | $F(1,7) = 0.033$, $p = 0.86$, $\eta^2_G = 0.0047$ |
| ITPC properties | | | |
| Precue-to-T1 slope (ΔITPC/s) | Mean = 0.052 SD = 0.047 | Mean = 0.048 SD = 0.053 | $F(1,7) = 0.60$, $p = 0.46$, $\eta^2_G = 0.079$ |
| Baseline ITPC (a.u.) | Mean = 0.26 SD = 0.13 | Mean = 0.32 SD = 0.11 | $F(1,7) = 0.51$, $p = 0.50$, $\eta^2_G = 0.068$ |
| ITPC T1 peak time (ms after precue) | Mean = 1201.75 SD = 3.38 | Mean = 1167.6 SD = 29.55 | $F(1,7) = 4.33$, $p = 0.076$, $\eta^2_G = 0.38$ |
| ITPC T1 normalized peak magnitude (a.u.) | Mean = 0.071 SD = 0.11 | Mean = 0.074 SD = 0.068 | $F(1,7) = 0.0033$, $p = 0.96$, $\eta^2_G = 0.00047$ |
| ITPC T2 peak time (ms after precue) | Mean = 1491.5 SD = 25.66 | Mean = 1468.2 SD = 29.8 | $F(1,7) = 1.26$, $p = 0.30$, $\eta^2_G = 0.15$ |
| ITPC T2 normalized peak magnitude (a.u.) | Mean = 0.064 SD = 0.10 | Mean = 0.087 SD = 0.071 | $F(1,7) = 0.15$, $p = 0.71$, $\eta^2_G = 0.021$ |

### Supplementary Figures

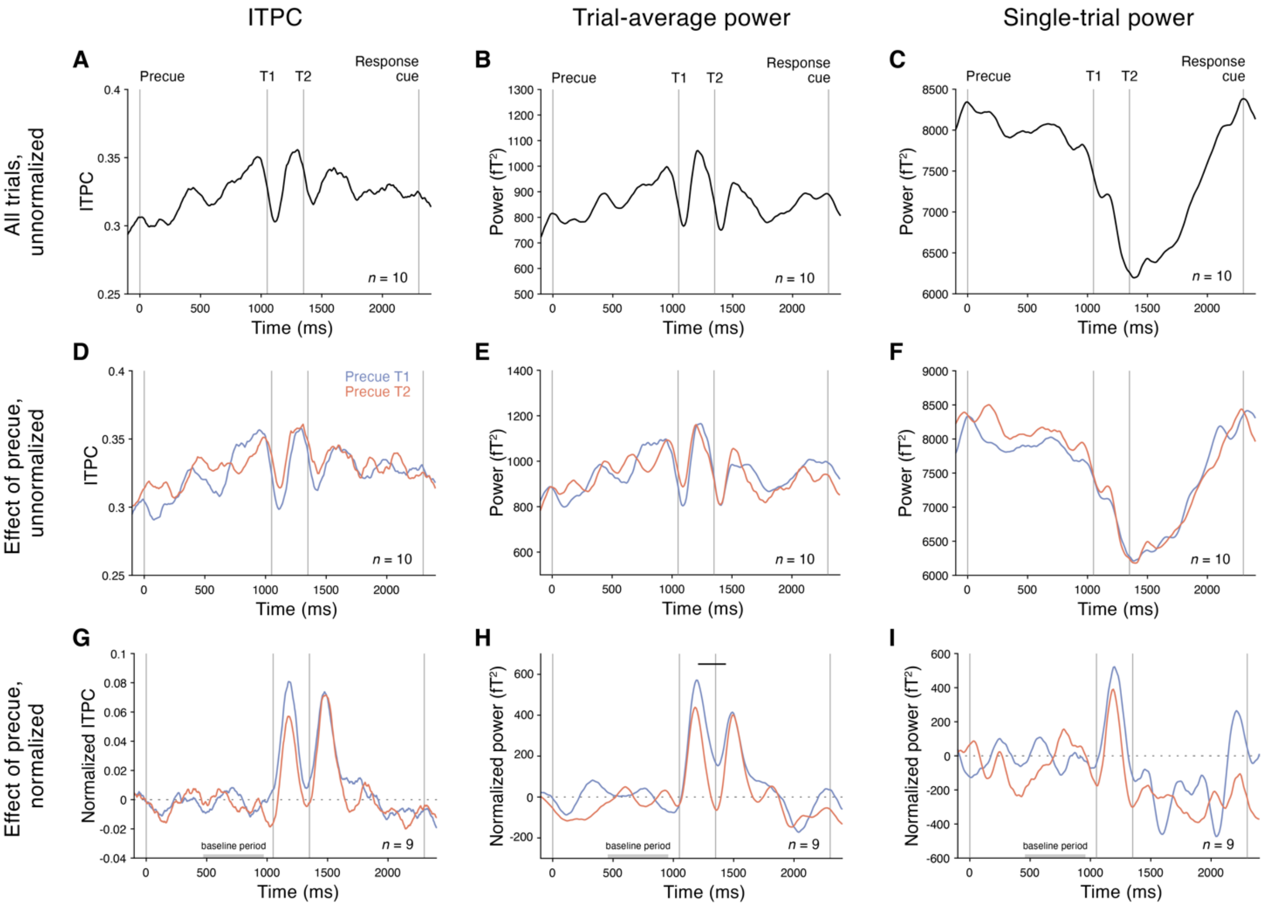


**Supplementary Figure 1**. Time series of all SSVER measures: ITPC (first column), trial-average power (second column), and single-trial power (third column). Unnormalized time series reveal pre-target anticipatory effects across all trials (first row) or as a function of the precue (second row). Time series normalized with respect to individual observer peak direction reveal post-target effects of the precue (third row). Horizontal bar indicates significant differences between precue conditions as shown by permutation tests comparing precue T1 vs. precue T2 conditions.

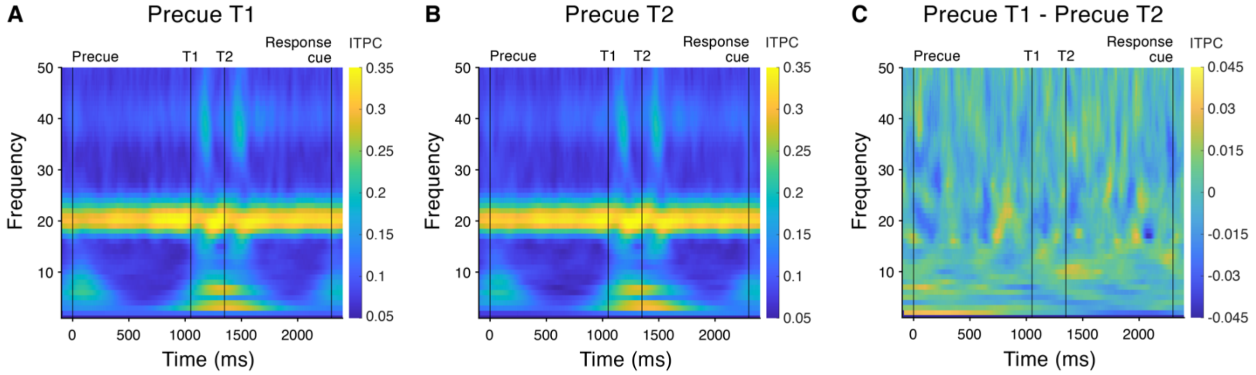


**Supplementary Figure 2.** Unnormalized ITPC spectra (*n* = 10) reveal no significant differences between precue types, as assessed using cluster-based permutation paired t-tests between (**A**) precue T1 and (**B**) precue T2 trials for each time (-100 to 2400 ms) and frequency (1-50 Hz) bin. (**C**) The difference of precue T1 and precue T2 spectra shows no significant clusters, indicating that ITPC for non-stimulus frequencies did not depend on temporal attention. Note that no significant difference is seen in the 20 Hz band of the difference spectrum because of observer based differences in the direction of target-evoked 20 Hz ITPC responses (see main text). In other frequency bands, target-evoked ITPC responses were only in the positive direction, making it appropriate to examine the unnormalized power spectrum for non-stimulus frequencies.

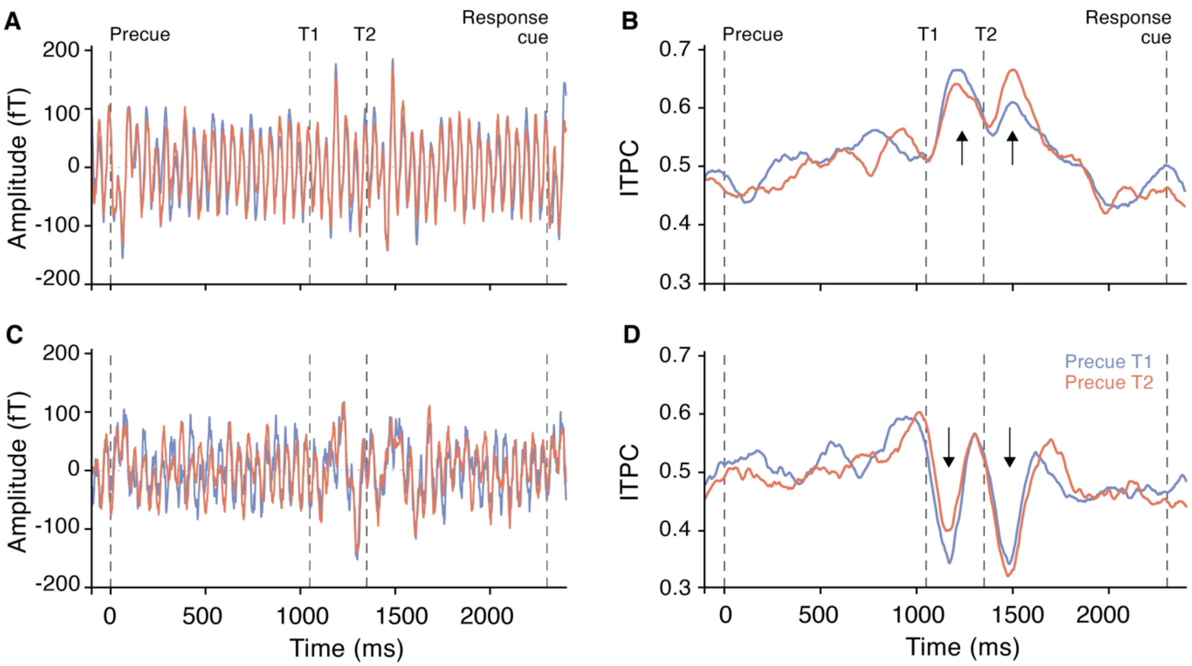


**Supplementary Figure 3.** ITPC time series and evoked ITPC response peaks for example observers. (**A,C**) Evoked time series and (**B,D**) 20 Hz ITPC time series for two example observers. Target-evoked ITPC responses increased for some observers (e.g., top row) and decreased for others (e.g., bottom row). Blue = precue T1, red = precue T2. Arrows show the directions of the target-evoked ITPC responses.

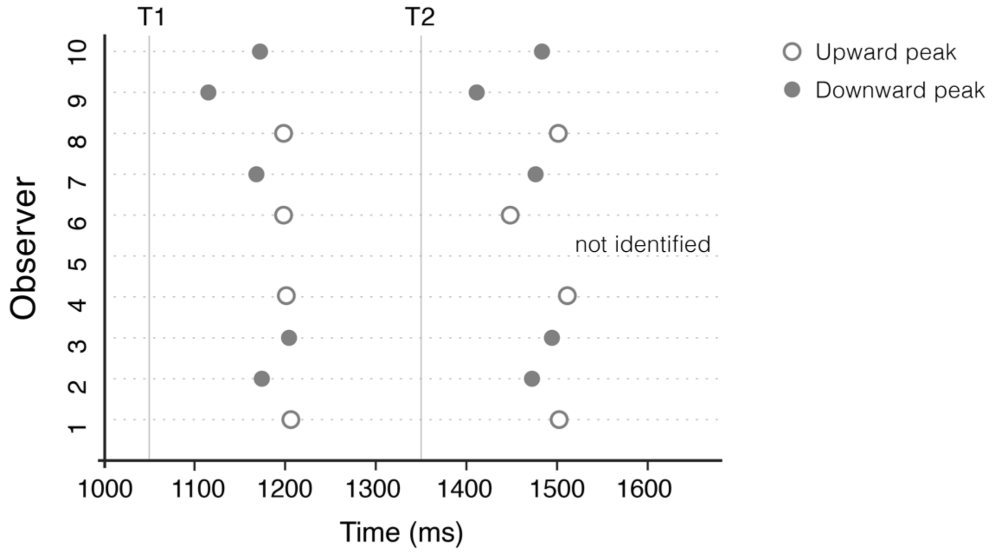


**Supplementary Figure 4.** ITPC peak times. The peak times for the target-evoked ITPC responses were consistent across observers, for T1 and T2. The directionality of the target-evoked ITPC responses was consistent within observers but varied across observers. Following the targets, the ITPC response increased for 4 of the 10 observers, decreased for 5 observers, and was unidentifiable for 1 observer.

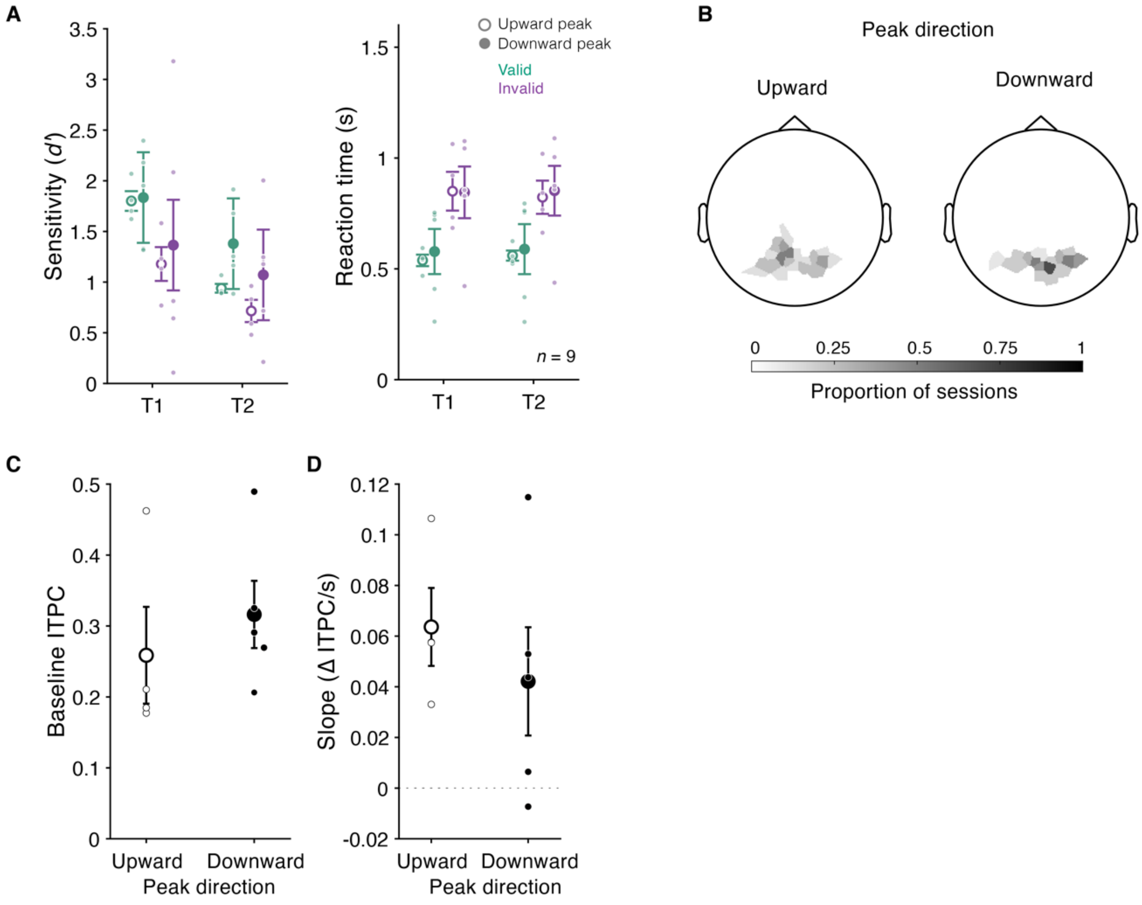


**Supplementary Figure 5.** Evoked ITPC response direction. The observer-specific direction of target-evoked ITPC responses was not predicted by (**A**) task performance: perceptual sensitivity (*d'*) and reaction time (s), (**B**) topographies of selected channels with highest SSVER power, (**C**) baseline ITPC, or (**D**) ITPC precue-to-T1 slope. Error bars indicate ± 1 SEM. Open circles indicate observers with upward target-evoked ITPC responses (n = 4), closed circles indicate observers with downward target-evoked ITPC responses (n = 5).
